# Multi-Modal Mass Spectrometry Identifies a Conserved Protective Epitope in *S. pyogenes* Streptolysin O

**DOI:** 10.1101/2023.12.02.569700

**Authors:** Di Tang, Carlos Gueto-Tettay, Elisabeth Hjortswang, Joel Ströbaek, Simon Ekström, Lotta Happonen, Lars Malmström, Johan Malmström

## Abstract

An important element of antibody-guided vaccine design is the use of neutralizing/opsonic monoclonal antibodies to define protective epitopes in their native three-dimensional conformation. Here, we demonstrate a multi-modal mass spectrometry-based strategy for in-depth characterization of antigen-antibody complexes to enable the identification of protective epitopes using the cytolytic exotoxin Streptolysin O (SLO) from *Streptococcus pyogenes* as a showcase. We first discovered a monoclonal antibody with an undisclosed sequence capable of neutralizing SLO-mediated cytolysis. The amino acid sequence of both the antibody light and the heavy chain was determined using mass spectrometry-based *de novo* sequencing, followed by chemical crosslinking mass spectrometry to generate distance constraints between the antibody fragment antigen-binding region and SLO. Subsequent integrative computational modeling revealed a discontinuous epitope located in Domain 3 of SLO that was experimentally validated by hydrogen-deuterium exchange mass spectrometry and reverse-engineering of the targeted epitope. The results show that the antibody inhibits SLO-mediated cytolysis by binding to a discontinuous epitope in Domain 3, likely preventing oligomerization and subsequent secondary structure changes critical for pore-formation. The epitope is highly conserved across >98% of the characterized *S. pyogenes* isolates, making it an attractive target for antibody-based therapy and vaccine design against severe streptococcal infections.

## Introduction

Antibody-guided vaccine design has emerged as a promising strategy to develop vaccines, which has shown promising results for viruses such as influenza (1) and HIV (2). The approach relies on molecular information from the adaptive immune response that is harnessed to design vaccine candidates that can elicit antibody responses of high magnitude and specific activity (3). The recent advances in the characterization of neutralizing antibody responses, together with new protein engineering methods, has catalyzed novel opportunities to rationally design next-generation vaccines (3). The neutralizing/opsonic monoclonal antibodies (mAbs) are used as guides to define the protective epitopes in their three-dimensional conformation, which requires detailed information of the atomic structure of the antigen–antibody complexes (4). Typically, structural biology methods such as nuclear magnetic resonance (NMR) spectroscopy, X-ray crystallography, and single-particle cryo-electron microscopy (cryoEM) provide insights into the 3-D structure of antigens and the antigen-antibody complexes (5). These methods are however, associated with high demands on sample quality and quantity or are limited by the molecular weight of the target antigens, impeding the identification of critically important epitopes in a high throughput manner.

Structural mass spectrometry is an emerging alternative to define epitopes of relevance for immunity. Hydrogen deuterium exchange mass spectrometry (HDX-MS) for example has proven useful for mapping epitopes between monoclonal antibodies and single antigens (6, 7). Additionally, recent studies have shown that epitope mapping can be accomplished by HDX-MS using polyclonal antibody mixtures, at least for small antigens (8, 9), without prior knowledge of the primary structure of the antibodies. HDX-MS can also reveal deprotected sites as a result of protein unfolding and allosteric effects (10), providing new information on protein dynamics in solution using limited amount of sample. Another approach is chemical crosslinking mass spectrometry (XL-MS), which facilitates the identification of proximal structural regions at the amino acid level (11). Protein samples are mixed with reagents that form covalent cross-links between defined residues in solution, and upon protease digestion, the resulting peptide pairs can be identified by tandem mass spectrometry (MS/MS) (12). The cross-linked peptide pairs generate distance constraints that can in conjunction with protein structural modeling resolve protein binding interfaces within, for example, antigen–antibody complexes (13, 14). This method is relatively fast, does not require complicated sample preparation protocols, necessitates relatively small amounts of starting material, can be applied to most proteins and is notably scalable to more complex mixtures of proteins (15). However, the primary structure information of both antigens and antibodies of interest is required.

*Streptococcus pyogenes* or Group A streptococcus (GAS) is a significant human pathogen responsible for considerable morbidity and mortality worldwide (16). Streptolysin O (SLO) is a 60 kDa pore-forming toxin ubiquitously produced by GAS, and belongs to a superfamily of pore-forming toxins as cholesterol-dependent cytolysins (CDCs). The first 69 N-terminal residues of SLO form a disordered region, followed by 3 discontinued Domains (D1-D3) and a membrane-binding Domain (D4) (17). The primary function of SLO is to bind cholesterol-rich membranes and induce cytolysis of eukaryotic cells. This depends on a multi-stage process that starts with SLO binding to cholesterol-rich membranes, followed by oligomerization to form a pre-pore and a conformational shift in Domain 3 to penetrate the membrane and induce cytolysis (18). In addition to this main biological function, SLO can also act as an immune-modulatory protein for neutrophils and impairs phagocytic clearance of GAS. Sub-cytotoxic levels of SLO have been found to suppress neutrophil oxidative bursts, enabling GAS to resist cell killing (19). Immunization with SLO can counteract this inhibitory effect, although the protection in certain challenge models falls short compared to M protein immunization (20). Moreover, SLO-deficient GAS strains exhibit decreased virulence in mouse models (21), while serum from SLO- or SLO toxoid-immunized mice can protect naïve mice against challenges with wild-type strains (17).

SLO is highly conserved and found in more than 98% of all characterized *S. pyogenes* isolates across the world (22), positioning SLO as a promising vaccine candidate. In fact, SLO is included as one of the antigens in at least three multivalent vaccines under investigation in pre-clinical and clinical trials (23). Current SLO-based toxoid vaccines have been developed by introducing point amino acid substitutions primarily in the membrane-binding loop of Domain 4 to prevent cytolysis during immunization (17, 24). Unlike the M protein, there are no reports to date of protective monoclonal antibodies with known primary structures against SLO, which impedes the identification of the protective epitope(s). In contrast, several cytolysis-inhibiting mAbs have been well characterized for pneumolysin O, a homologous CDC secreted by *Streptococcus pneumoniae*, however, the epitope mapping was mainly conducted using overlapping peptide fragments and domains (25).

In this study, we propose a novel multi-modal mass spectrometry strategy and sequenced a neutralizing monoclonal antibody to determine the protective epitope by investigating SLO-antibody complex in its native state. The epitope is located in Domain 3 and is conserved across >98% of the GAS genomes. The findings also showcase multi-modal mass spectrometry as a versatile and promising strategy for identifying protective epitopes within bacterial antigens.

## Results

### *De novo* sequencing and modelling of a neutralizing monoclonal antibody

In our proposed multi-modal mass spectrometry workflow, epitopes targeted by monoclonal antibodies (mAb) are identified by first using *de novo* mass spectrometry sequencing to determine the primary structure of mAb, and then, XL-MS to generate distance constraints within the antigen-antibody complex. In the next step, the epitopes are defined by an information-driven docking protocol to predict the most important residues in the binding interface between the fragment antigen-binding (Fab) region and the antigen that can be experimentally validated by HDX-MS and reverse-engineering of the derived epitope (**Fig. 1A-B**).

**Fig. 1.**
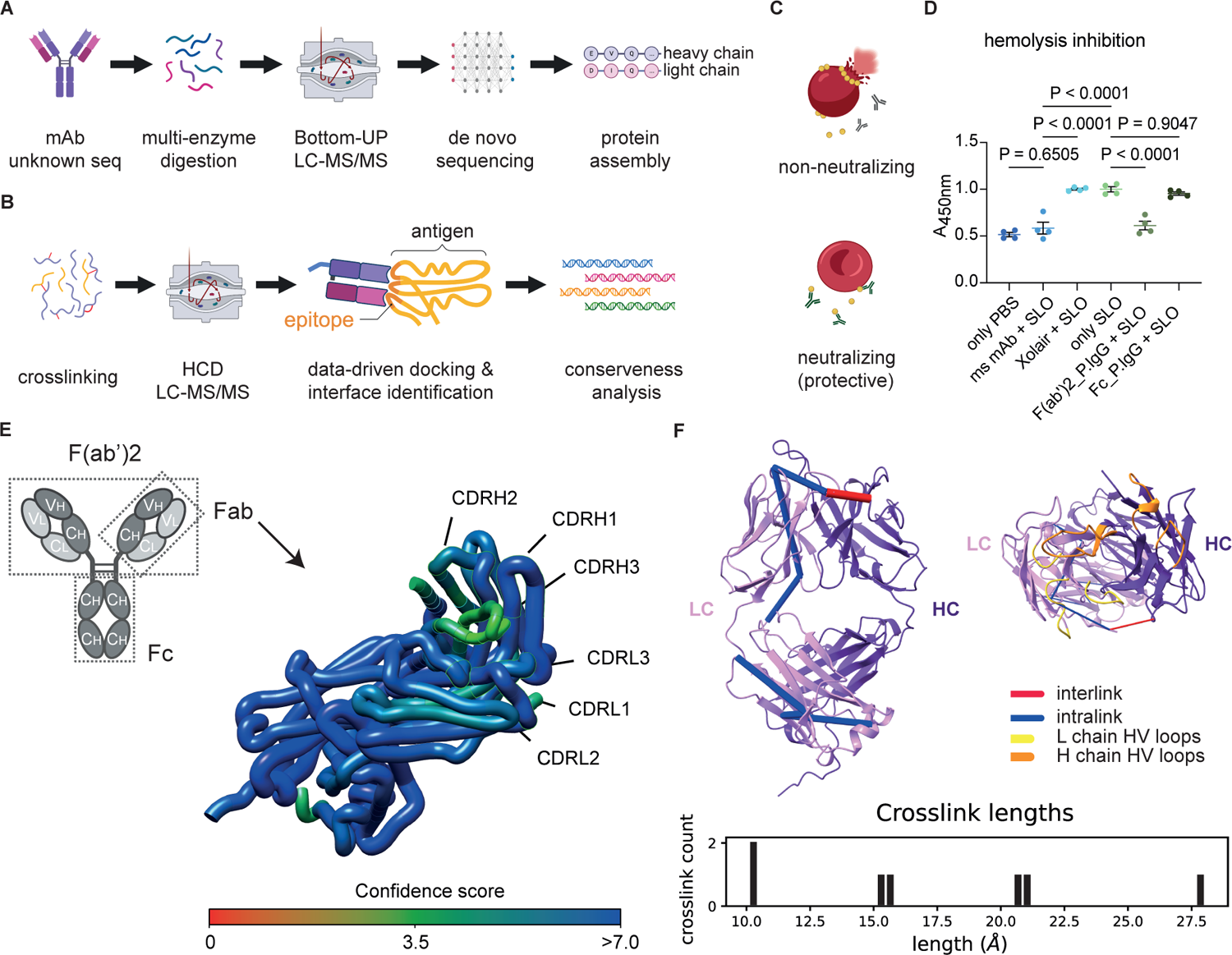
**A-B).** Graphical illustration of the workflow outlining *de novo* protein sequencing of the monoclonal antibody (mAb) using multi-enzyme digestion, bottom-up mass spectrometry *de novo* sequencing (BU-MS), and sequence assembly, followed by crosslinking mass spectrometry (XL-MS) to define XL distance constraints for exploratory modelling, pairwise antigen-antibody complex docking, and assessment of residues involved in the binding interface. **C).** Schematic representation of the inhibition assay used to select neutralizing antibodies (nAb). **D).** Hemolysis inhibition was determined by co-incubating different antibody samples with wild-type SLO together with diluted sheep red blood cells. The antibody-mediated inhibition of hemolysis was quantified by measuring the amount of released hemoglobin in the supernatant (ms = a murine IgG1, Xolair = an anti-IgE IgG1 mAb, P.IgG = isolated IgG from convalescent plasma of a single donor recovering from a recent GAS infection). P values from ANOVA group analysis are indicated. **E).** Schematic illustration of an IgG1 antibody, illustrating two heterodimeric chains comprised of both variable (VH and VL) and constant regions (CH and CL). A tertiary model generated from AlphaFold-Multimer of the *de novo* sequenced Fab domain, with the confidence score for each residue shown by the color gradient. **F).** To validate the model, XL-MS was applied to identify inter/intra-nAb crosslinks. Inter-protein (red) and intra-protein (blue) crosslinks are displayed as pseudo bonds within the predicted Fab structure. The bar plot in the bottom shows all identified crosslinks, with the y-axis representing the count and the x-axis representing Cα-Cα distances. Light chain (LC) and heavy chain (HC) are distinguished by color codes, and the proABC-2-predicted-hypervariable (HV) loop regions are highlighted by yellow, or orange as indicated by the color legends.

Antibody protection can be divided into neutralization and/or opsonization. In this study, we defined antibody protection by the degree of neutralization of SLO-mediated cytolysis by antibodies (**Fig. 1C**). As a starting point, we screened readily available anti-SLO mAbs from public and commercial sources that could bind to SLO with high specificity leading to neutralization of SLO-mediated cytolysis. Using a sandwich ELISA, we identified a murine mAb that bound specifically to SLO (**Fig. S1**). Neutralization was further assessed by incubating different antibody samples with active SLO, followed by the addition of diluted red blood cells from sheep. Inhibition of cytolysis was quantified by measuring the reduction of released hemoglobin in the supernatant. The neutralizing mAb (nAb) completely inhibited the SLO-mediated cytolysis compared to the incubation with the unrelated murine IgG1 Xolair (**Fig. 1D**, **Table S1**). To further investigate if neutralizing antibodies occur after bacterial infection, we isolated IgG from plasma of a single donor recovering from a recent GAS infection. By separately comparing the neutralizing effect of the F(ab’)2- and Fc-fragments, we could demonstrate that neutralizing antibodies can arise after a GAS infection and that neutralization is mediated by the Fab-domain (**Fig. 1D**; **Fig. 1E**, top left).

To obtain the full amino acid sequence of the nAb, we digested the antibody using four different proteases with different cleavage specificity and used a combined *de novo* sequencing strategy based on three different search engines to sequence proteolytically overlapping nAb peptide fragments. The overlapping peptide fragments were assembled into a high-confidence full-length sequence including the complementary determining regions (CDRs) involved in the antigen binding for both the light and heavy chains followed by modeling of the tertiary structure of the Fab-domain (**Fig. 1E**, right). To validate the model, we used AlphaFold-Multimer (26) to predict the Fab structure (ipTM + pTM = 0.848; inter-chain pDockQ = 0.688) followed by XL-MS to generate intra- and inter-molecular distance constraints within and between the nAb heavy chain (HC) and the light chain (LC). The identified interlinks and intralinks (**Table S2**) conformed to the modeled Fab structure without violating the Cα-Cα upper distance limit of 30 Å (**Fig. 1F**, top left and bottom). Furthermore, prediction of the antibody residues and type of interactions involved in the intermolecular interaction with antigens were determined by proABC-2 (27). This analysis pinpointed residues 26-32, 50-52, and 91-96 in the light chain as hypervariable (HV) contact loop region and the residues 26-31, 52-57 and 100-114 in the heavy chain as HV contact loop region (**Fig. 1F**, top right). Collectively, the identified intramolecular distance constraints show that the primary structure together with the modeled Fab structure is of sufficient quality to infer the epitope via XL-MS and information-driven docking.

### Predicting the binding sites by XL-MS and inferring epitope by interaction analysis

To identify the epitope targeted by the nAb, we incubated SLO at a 1:1: molar ratio with the nAb followed by chemical crosslinking using disuccinimidyl glutarate (DSG) or disuccinimidyl suberate (DSS). Subsequent mass spectrometry analysis identified five cross-linked peptide pairs between SLO-Fab/LC and three peptide pairs between SLO-Fab/HC. The DSG and DSS cross-linkers have different lengths of the spacer arm, resulting in partially overlapping peptide pairs. Three of the cross-linked peptide pairs were identified by several independent crosslinked spectrum matches (CSM) for both DSS and DSG (**Fig. 2A-B**, **Table S3-4**) and representative MS/MS spectra for the two most frequently observed crosslinked peptide pairs (SLO/K375-FabLC/K31 and SLO/K403-FabHC/K62) are shown in **Figure 2C-D**.

**Fig. 2.**
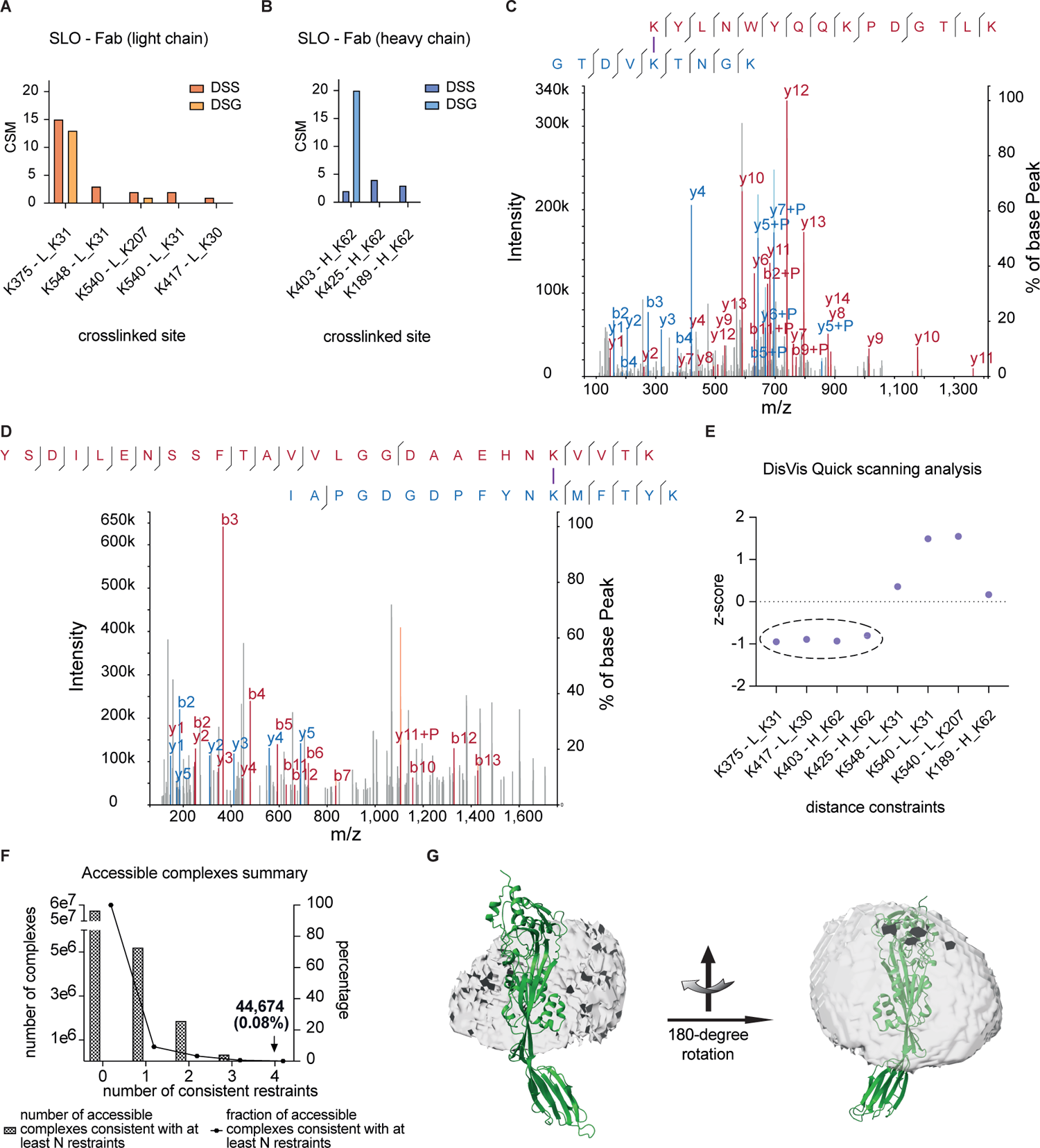
**A-B).** The number of cross-link spectrum matches (CSM) for the identified inter-protein DSG- and DSS-crosslinked peptide pairs between SLO-Fab light chain and SLO-Fab heavy chain, K, lysine; L, light chain; H, heavy chain. **C-D).** Representative MS/MS spectra of the two predominant crosslinked peptides are illustrated with annotated fragment ions. The top panel and lower panel represent the SLO/K376-FabLC/K31 and the SLO/K403-FabHC/K62 crosslinks, respectively. **E).** DisVis Quick scanning analysis was used to filter experimentally derived inter-protein distance constraints for Fab-SLO modelling. Each DSS-constraints are plotted on the x-axis with the z-score value on the y-axis. The dotted circle indicates the group of 4 constraints with low z-scores that was used for downstream analysis. **F).** Histogram showing the reduction of accessible complexes based on the number distance constraints. The remaining 44,674 complexes consistent with all 4 distance constraints was used for downstream analysis. **G).** The accessible interaction space formed by the filtered 44,674 complexes shown as a shell centered around Domain 3 in SLO.

To select the most informative distance constraints, DisVis (28) quick scanning analysis was applied to the identified crosslinks (XLs). Four of the XL distance constraints displayed a negative z-score, an indicator of relevance, and were selected for further analysis. The two XL constraints with the highest number of CSMs were included among these four selected XL distance constraints (**Fig. 2E**, **Table S5**). In the next step, the filtered XL distance constraints were used to guide the complete DisVis interaction analysis, to identify the putative residues and to infer the binding interface (**Fig. S2**). By setting SLO as the fixed chain and Fab as the scanning chain without using any XL distance constraints, nearly six million conformations were generated as an initial sampling space. This large conformational space was produced by allowing the Fab chain to freely translate and rotate relative to the fixed SLO chain. All surface-exposed residues on both interacting partners, with a relevant solvent accessible (RSA) value greater than 40% (**Table S6**), were considered potential interactive residues. This approach was aimed to provide a complete representation of all possible interaction scenarios between SLO and the Fab fragment. The stepwise inclusion of the four most informative cross-links reduced the conformational space by removing conformations that violated the introduced distance constraints, reducing the number of conformations to a final set of 44,674 (0.08%) when including all four XL distance constraints (**Fig. 2F**, **Table S7**). The final interaction space was produced from all possible SLO-Fab complexes using the filtered 44,674 conformations indicates that the epitope resides within Domain 3 of SLO (**Fig. 2G**). Finally, we used interaction fraction (IF) index (28) to predict the residues of importance for the antigen-antibody binding interface. On the Fab side, the most important residues with an IF index higher than 0.5 were located in the CDRs for both the heavy and the light chain (**Fig. 3A**, **Fig. S3**). Reciprocally, the most important residues in SLO were located in two distinct discontinuous regions between amino acids 187-244 and 357-440 with the latter region being the most significant one based on the IF indexes (**Fig. 3B**, **Fig. S3**). Highlighting these regions in the crystal structure of SLO (PDB: 4HSC) shows that the epitope is discontinuous and located within Domain 3 and composed of 3 alpha-helices and 2 beta-sheets (**Fig. 3C**).

**Fig. 3.**
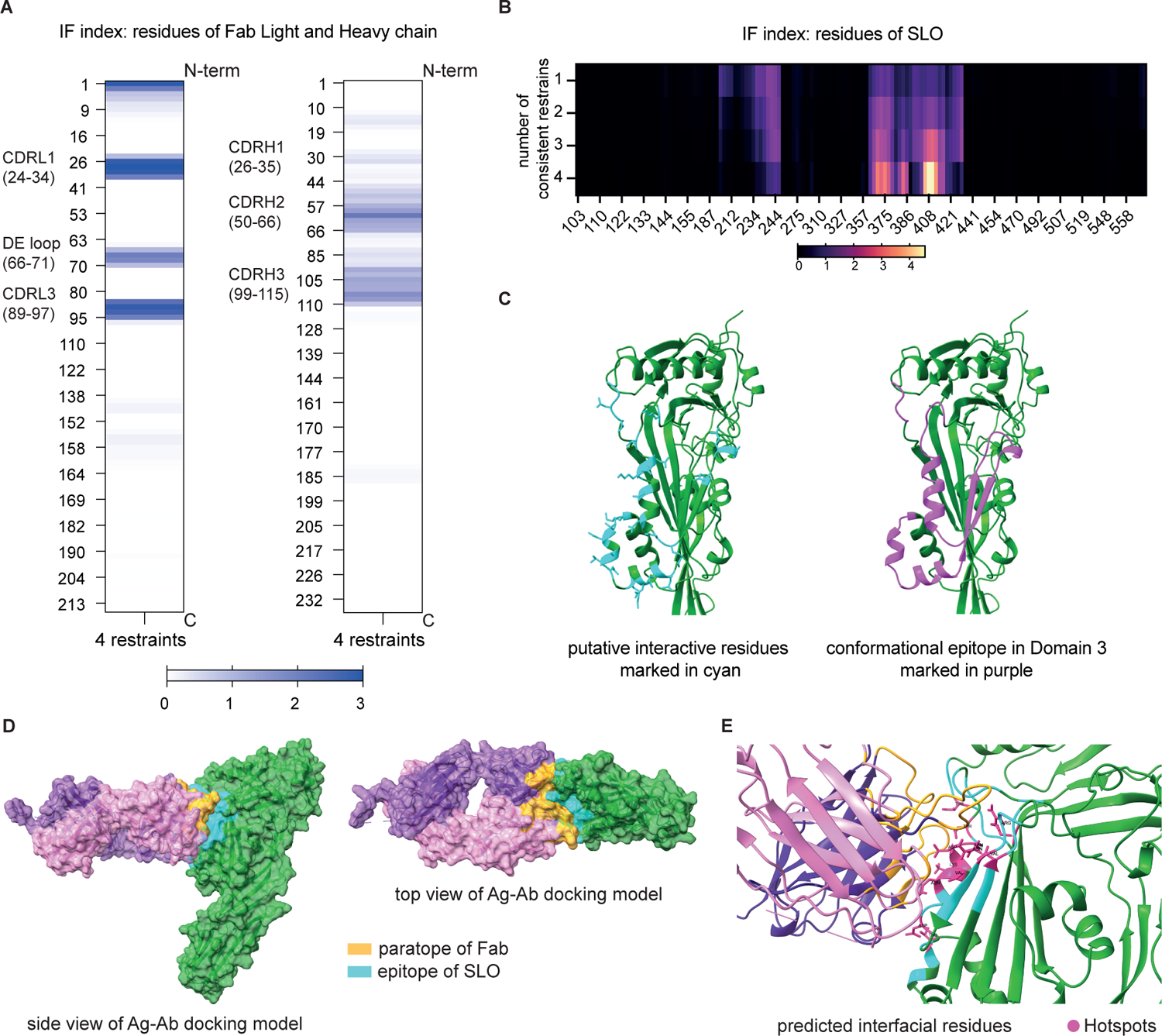
**A).** Heat maps displaying the interaction fraction (IF) score of each residue from the light and the heavy chain respectively, with a color bar with the IF index ranging from 0 to 3. **B).** Heatmap showing the IF index of all accessible SLO residue based on the number of distance constraints used. Color bar shows IF value from 0 to 4.6. **C).** Left, display of most likely interacting residues (with an IF >0.5) with sidechains shown as sticks on the tertiary structure colored in cyan. Right, overview of conformational epitope harboring the indicated residues in the SLO 3D crystal structure, colored in purple. **D).** Top-ranking model generated by HADDOCK 2.4 antigen-antibody docking protocol with a loose XL-MS derived epitope definition, predicted paratopes as hypervariable (HV) loop contacts and the two most frequently observed distance constraints as input. The PRODIGY-predicted epitope and paratope from the top-ranked complex are color-coded correspondingly. **E).** Interface residues were identified and classified from the top-ranked Fab-SLO complex model, with hotspots highlighted in hotpink and labeled with residue name to represent residues predicted to contribute most significantly to the intermolecular interaction.

### Pinpointing interface by distance-information-driven docking of the Fab-SLO complex

The DisVis analysis, the IF-index along with acquired XL distance constraints enabled us to predict a loosely defined epitope located in Domain 3 of SLO. To further determine the interaction site, we used the HADDOCK 2.4 antigen-antibody docking protocol (29, 30). This protocol uses the interface information from DisVis to generate more accurate starting conformations for iterative docking and refinement of pairwise complexes. We tested different information inputs such as no epitope information, loose epitope definition, loose epitope definition + XL distance constraints, to guide the docking process and found that the DisVis-derived loose epitope combined with the two most frequently observed XL constraints (SLO/K375-FabLC/K31 and SLO/K403-FabHC/K62) as the center of mass, resulted in the most significant cluster comprised of 240 of the 400 models (**SI text**). The top-ranked Fab-SLO complex model from this cluster holds a combined HADDOCK score of −145 (**SI text**), where normally a HADDOCK combined score of <-120 is considered a confident score (29–31). The selected Fab-SLO complex is depicted from two perspectives, highlighting the paratope and epitope composed of interfacial residues as predicted by PRODIGY (32). Next, the SpotOn algorithm (33), which predicts possible sidechain interaction between pairs of residues within pairwise protein complex, was applied to the top modeled Fab-SLO complex structure to identify the interacting residues, referred to as hotspots. This analysis indicates that residues Arg346, Val393, Glu400, Asn402, Thr406, Asp410, Val411, Asn414 and Asp418 in SLO are involved in the protein binding interface (**Fig. 3E**, **Table S8**).

### HDX-MS validation of the epitopes derived by XL-MS and information-driven docking

To further validate the results from the distance-information-driven docking, we used HDX-MS to investigate changes in deuterium uptake in SLO residues when bound to the nAb and to further investigate any changes in SLO protein dynamics upon binding. The HDX-MS experiments generated high-confidence annotation of 202 peptides corresponding to a sequence coverage of 95.7% with a high degree of overlap (**Fig. 4A**). The changes in deuterium uptake (ΔDU) over four deuteration intervals (0s, 60s, 1800s, and 9000s) for each identified peptide were aggregated and shown in a butterfly plot in **Figure 4B**. Overall, the changes of deuterium uptake between SLO alone (apoSLO) and SLO in complex (nAb-SLO) were minor, except for the peptides starting from Thr390 to Phe409. Statistical testing identified the peptides spanning 392-421, 398-409, 392-422, 391-408, 390-433, 392-409, 390-408, 390-409, 390-410 and 390-422 to be significantly protected upon binding of the nAb to SLO (**Fig. 4C**). The kinetic plots in **Figure 4D** illustrate deuterium uptake from two representative peptides across the four deuteration intervals, with the level of significance indicating protection exerted by the nAb. The protected peptides were all found in Domain 3 of SLO (**Fig. 4E**), and near-perfectly overlap with the epitopes predicted from the XL-MS interaction analysis and the Fab-SLO complex docking model (**Fig. 4F**). Importantly, the HDX-MS results further show that SLO does not undergo major conformational change upon nAb binding.

**Fig. 4.**
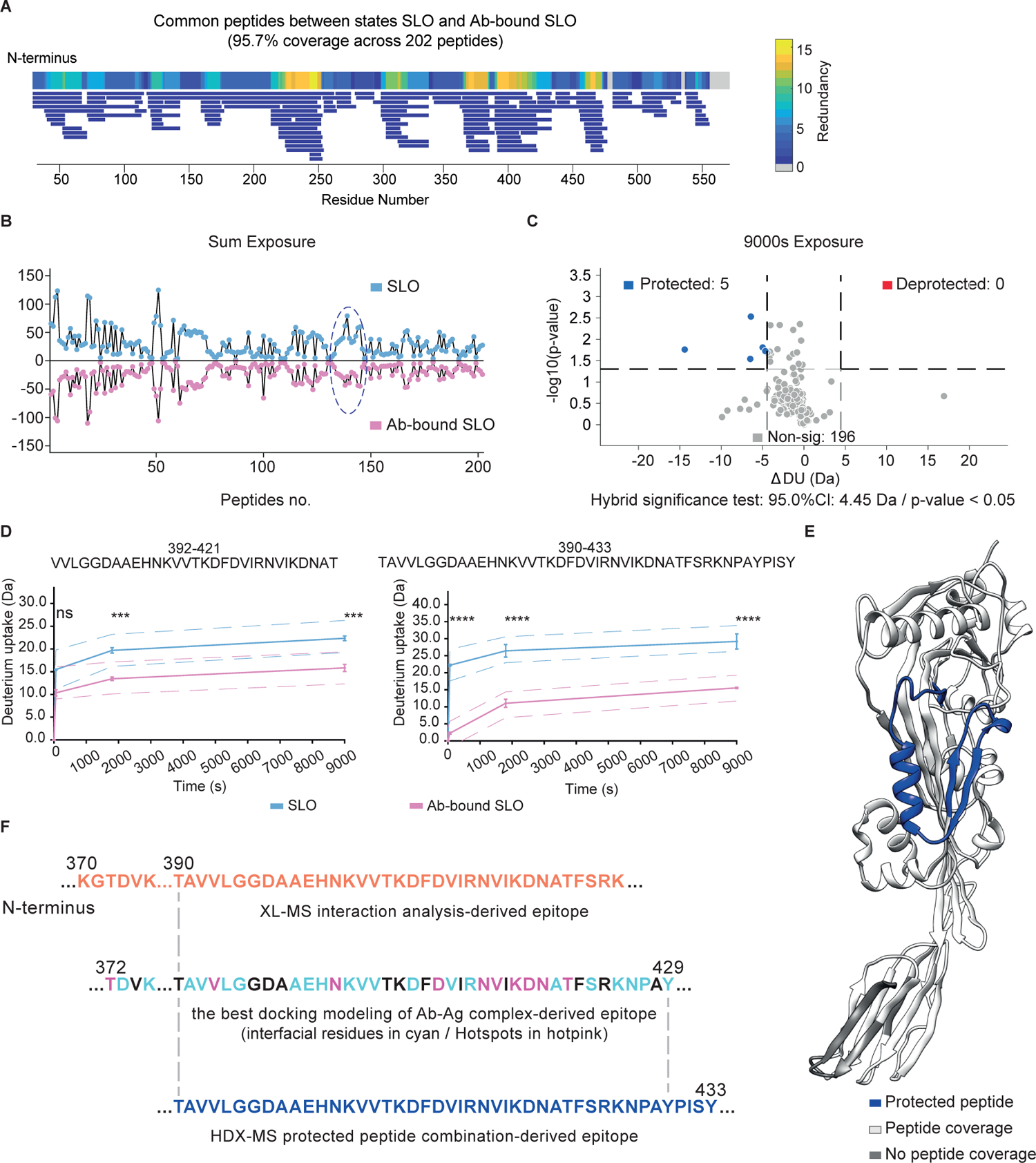
**A).** The redundancy map of 202 common peptides identified in both SLO and nAb-SLO complex from the HDX-MS experiments. Peptide fragments are displayed as bar with the length correlating to peptide length, with residue redundancy visualized by a color gradient from 0 to 16. **B).** The cumulative Da changes due to deuterium uptake across all peptides are shown in a butterfly plot in the top, which compares the peptides found in both SLO and Ab-bound SLO. **C).** A volcano plot in the bottom presents significantly protected and deprotected peptides, with thresholds set at |ΔDU| >4.45 Da and a p-value of <0.05. **D).** Kinetic plots displaying two representative protected peptides of SLO under Ab binding at 0s, 60s, 1800s, and 9000s deuteration intervals. Thresholds were globally calculated with ‘ns’ as non-significant, ‘***’ as ≥99% significance and ‘****’ as ≥99.9% significance. **E).** The 3D model of SLO displaying the protected peptides after 9000s deuteration. Here, residues from the protected peptides are highlighted in blue, while residues not detected in the HDX-MS analysis are depicted in dark grey. **F).** Comparison between the three protective epitopes derived from XL-MS, information-driven docking and HDX-MS, highlighted in colors from Figure 3B, Figure 3E and Figure 4F, with residue position number on top of the sequence.

### Validation of the epitope by a novel reverse-engineered construct

In the final experiment, we designed and produced a new protein construct referred to as d3_m, to verify that the identified discontinuous epitope was sufficient for nAb binding. Here, we used in-silico stabilization analysis to merge the two non-continuous regions of domain 3 that were identified by XL-MS and HDX-MS by replacing the stretched region from Lys298-Arg346 with a short linker sequence of four amino acids (Ala-Pro-Asn-Gly). The entire d3_m construct is 128 amino acids long and is 76.3% shorter than the full length SLO. Initial structure prediction by AlphaFold2 (34) indicated that the design is in close alignment with the SLO crystal structure with a root-mean-square-deviation (RMSD) of 0.715 Å when comparing the structured regions (**Fig. 5A**). To determine whether the nAb could bind to the construct, we expressed and purified d3_m in *E.coli* and evaluated the binding using ELISA. The nAb binds specifically to the d3_m in an equal manner compared to intact SLO, while an unrelated IgG (Xolair) showed no binding (**Fig. 5B**, **Table S9**). Furthermore, the equilibrium dissociation constant as Kd determined by non-linear regression analysis, indicates that the nAb binds marginally stronger to intact SLO compared with the binding to d3_m (**Fig. 5C**, **Table S10**).

**Fig. 5.**
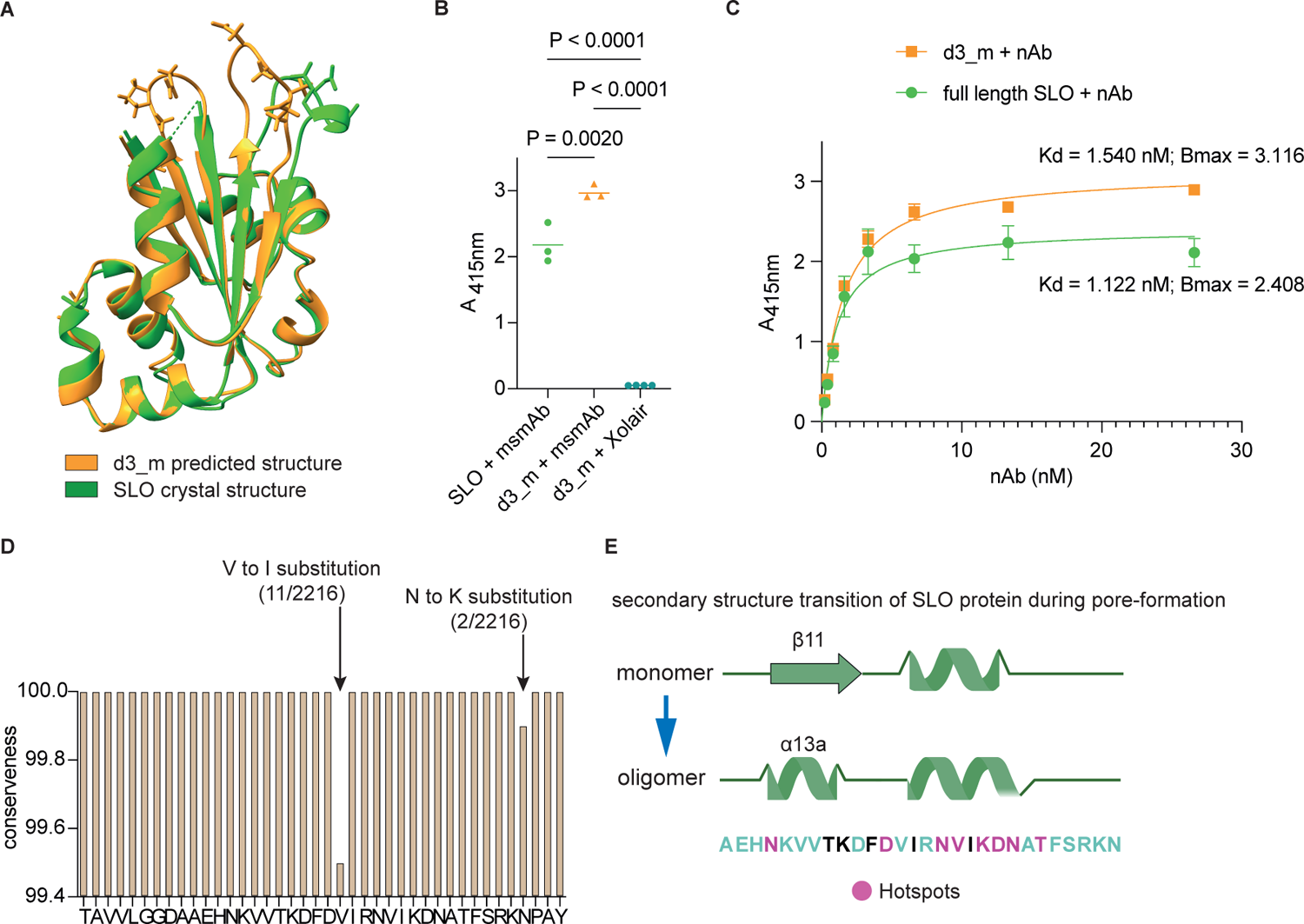
**A).** Structure superimposition and alignment were performed on the predicted structure of the novel construct d3_m and the corresponding region extracted from the crystal structure of SLO (PDB ID: 4HSC). The visualization of side chain highlights the conformationally dissimilar loop regions. **B-C).** ELISA was conducted to assess the nAb specificity against the d3_m construct by one-way ANOVA group analysis, also to estimate the corresponding binding via the equilibrium dissociation constant (Kd) and maximum specific binding (Bmax), derived from the non-linear regression analysis. Adjusted P values are added accordingly. **D).** Level of conserveness of the epitope residues across 2216 available GAS genomes. The bar indicates the degree of conserveness for the residues that constitute the converged epitope. **E).** Streptolysin O, similar to other cholesterol-dependent-cytolysins, undergoes conformational changes, particularly, in the secondary structure of Domain 3. The nAb Fab-recognized epitope colored as Figure 4E overlaps a portion of these structurally critical motifs shown as green helix and sheet cartoons.

### High conserveness of the identified epitope and protection mechanism speculation

To investigate if the epitope is conserved in GAS, we first mined streptococcal genomes available in the public domain (35). Of the available 2216 GAS and 154 Streptococcus dysgalactiae genomes, 98% of the GAS and 56% of the *S. dysgalactiae* strains carry the SLO gene using a >98% sequence similarity compared to the SLO protein with UniProtID: P0DF96. In the next step, we assessed the level of conservation for the epitope by querying the epitope sequence in 2216 publicly available GAS genomes. Among all genomes that carried SLO, the epitope was nearly 100% conserved apart for 0.6% of the SLO sequences having a semi-conserved amino acid point substitution from valine to isoleucine, and/or asparagine to lysine (**Fig. 5D**). Like other CDC family members, streptolysin O forms a pore on cholesterol-rich membrane through a multi-stage process that includes initial binding to the cellular membrane followed by oligomerization. During this process Domain 3 undergoes a significant conformational change including transformation of the D3 central beta sheet 11 to alpha helix 13 where hotspot Asn402 is located (**Fig. 5E**). These secondary structure shift triggers oligomerization, pre-pore formation and final transition into a pore. Both the HDX-MS and XL-MS results show that the nAb binds precisely to this critical region responsible for oligomerization, hence possibly preventing the secondary structure shift required for the subsequent pore-formation.

## Discussion

Monoclonal antibodies play an increasingly important role for the definition of protective epitopes that can serve as a starting point for the rational design of next-generation vaccine candidates (36). However, the availability of protective monoclonal antibodies against antigens from bacterial pathogens remains scarce, and their use in defining protective epitopes poses a considerable analytical challenges. Here, we demonstrate the versatility of mass spectrometry-based proteomics by i) *de novo* sequencing a neutralizing monoclonal antibody that inhibits SLO-mediated cytolysis of eukaryotic cells and ii) using a combination of XL-MS, HDX-MS, and integrative computational modeling to identify protective epitopes using antigen-antibody complexes in their native state.

Recent developments in deep learning algorithms in structural biology, such as AlphaFold (26, 37, 38), have excelled in predicting protein-protein interactions. Still, modelling of antibody-antigen complexes remains challenging due to limited co-evolutionary features and the limited availability of antigen-antibody co-crystal structures (39). Experimentally derived distance constraints that define or infer the binding interface are of importance to guide the modelling of antibody-antigen complexes and to improve the accuracy of the deep learning algorithms (40). Over the years, several MS-based techniques have been developed capable of generating distance constraints for modelling such as XL-MS, HDX-MS, limited-proteolysis MS, and covalent labeling (40, 41). These MS-based techniques typically rely on low quantity of starting materials and have comparably fast sample preparation strategies while providing informative distance constraints (42), information regarding solvent accessibility and insights into conformational ensembles in solution (43) and protein dynamics (40, 44, 45) under native conditions (41). At the same time, these techniques are also associated with limitations. Although XL-MS can be applied to most proteins and is scalable to more complex protein mixtures, the method is limited to protein complexes where cross-linkable residues are in close proximity. For antibody-antigen complexes, XL-MS comes with the additional drawback that the primary structure of both antibody and antigen is required for searching crosslinked peptide pairs. In contrast, mapping epitopes with HDX-MS does not necessitate primary structure information of the antibodies. In addition, HDX-MS can reveal deprotected sites indicating protein unfolding and allosteric effects (10), providing new information of protein dynamics in solution. On the other hand, HDX-MS provides in most applications relatively low peptide-level structural resolution and requires high-sequence coverage of the proteins under examination, limiting the scalability of the method to complex protein mixtures. Previous studies have shown that implementation of several complementary methods can considerably enrich the structural information, compared to any stand-alone method (46, 47). In our study, HDX-MS was used to validate the computationally derived epitope mapping guided by XL-MS. In addition, HDX-MS can also be used to exclude XL-derived distance constraints. In our case, HDX-MS did not support several distance constraints identified by XL-MS (SLO/K548-FabLC/K31, SLO/K540-FabLC/K31, SLO/K540-FabLC/K207 and SLO/K189-FabHC/K62) and these were considered to be weak or transient interactions captured by XL-MS. This cross-validation underlines the use of integrative multi-modal mass spectrometry to decipher antibody-antigen interactions. Beyond confirming the epitope, HDX-MS was used to demonstrate that binding of the nAb to SLO did not introduce any major conformational shift, similar to the one occurring during pore formation (18). This emerging concept of integrating several MS-based techniques extends new opportunities to obtain structural information and protein dynamics for other generic protein-protein complexes, which require considerable effort to crystallize, or which do not crystallize at all.

Like most streptococcal antigens, there were no available protective monoclonal antibodies with known primary structure against SLO. Here, we solved this by *de novo* sequencing an existing monoclonal antibody and used the epitope information to unveil the protective mechanism exerted by the antibody. Interestingly, the epitope region is highly conserved across many genomes from both group A and Streptococcus dysgalactiae indicating that rationally designed vaccines based on this epitope could potentially confer cross-species protective protection against SLO produced from both species. Yet the epitope remains quite distinct from other CDC family proteins like pneumolysin O. This notion is supported by previous reports that have demonstrated the neutralizing monoclonal antibody used here is specific against SLO (48). On the other hand, there are currently a few neutralizing mAbs that bind specifically to pneumolysin O (25). An interesting future prospect would be to investigate the epitopes in pneumolysin O that are recognized by these monoclonal antibodies, and to compare them to the protective epitope of SLO identified from current study. Such information would provide basis for the future design of vaccine candidates that display epitopes specific for certain members from the CDC family or possibly against several, or of all the members. These findings could additionally guide future development of targeted antibody treatments for GAS-infected patients through passive immunity therapies.

*Streptococcus pyogenes* is a significant human pathogen responsible for considerable morbidity and mortality. There has been a long history of vaccine development against GAS. However, the path to a successful vaccine candidate has been impeded by scientific and regulatory barriers (23). Over the last five years, efforts within GAS vaccine development have been re-invigorated. Still, there are only eight early-stage vaccine candidates that have shown effect in preclinical models and so far, only one candidate has demonstrated safety and immunogenicity in a phase 1 trial. SLO is a notable vaccine candidate and one of the antigens in at least three multivalent vaccines under investigation in pre-clinical and clinical trials (23). Current SLO-based toxoid vaccines have been developed by introducing amino acid substitutions primarily in the membrane-binding loop of Domain 4 to prevent cytolysis during immunization (17, 24). However, the rational design of SLO-based vaccines against *S. pyogenes* has still not been demonstrated. In this context, our study provides a possible starting point for the design of a vaccine that would specifically present a highly conserved and protective epitope. The novel construct d3_m was designed and reversed-engineered based on epitope structure derived from our multi-modal mass spectrometry approach. Modelling of the construct showed close structural similarity with the native structure of SLO. Significantly, the nAb demonstrated binding affinity to the d3_m almost equivalent to its binding to the intact SLO, indicating that reverse-engineering of the epitope is feasible and successful. Higher Bmax (maximum specific binding) value of d3_m also demonstrated the better epitope accessibility for the nAb on this construct. In this context, a multivalent or multi-domain-derived vaccine design, integrated with epitopes targeted by existing or future neutralizing/protective antibodies, has the potential to enhance ongoing efforts in developing next-generation vaccine against GAS.

## Methods and Materials

### SLO protein and recombinant protein production

Active SLO protein was obtained commercially (Sigma Aldrich) or expressed and purified recombinantly in-house. The complete SLO sequence (Uniprot ID: P0DF96), excluding the signal peptide, was engineered to incorporate an N-terminal Strep-HA-His-tag. The plasmid carrying the tag-SLO gene was synthesized and assembled by the Lund University Protein Production Platform, subsequently transformed into BL21(DE3) Competent Cells (Thermo Scientific). The induction and purification of the target tag-SLO protein was carried out in-house following a previously published protocol (49). The purity and sequence of the tag-cleaved SLO protein were confirmed by SDS-PAGE (Bio-Rad) and bottom-up MS (BU-MS). The reverse-engineered d3_m construct was produced and purified by the Protein Production Sweden Umeå node.

### nAb specificity and neutralization against SLO

The murine nAb was from Abcam (AB23501, Lot No: GR3430639). For ELISA assays, an equivalent quantity of SLO protein or the d3_m construct (ca. 3 ug) was immobilized onto a MaxiSorp plate (Thermo Scientific), which was subsequently incubated with either 0.4 ug nAb (followed by serial half-dilution), Xolair or 1xPBS (Phosphate buffer saline tablet, Sigma Aldrich) as a background control. Post thorough washing, a secondary HRP-conjugated anti-mouse IgG goat antibody (Bio-Rad) and HRP substrate kit (Bio-Rad) were added in sequence. After 3 min development, the plate was read in a microplate reader (BMG Labtech) at a wavelength of 415 nm. To compare binding of the nAb against full length SLO and d3_m construct, Prism 10 built-in non-linear regression (Equation: One site -- Specific binding) mode was applied to derive the Kd and Bmax with 95% confidence interval. For the SLO cytolysis inhibition assay, sheep blood cells (Thermo Scientific, Oxoid) were first diluted with 1xPBS, followed by the addition of 100 ng of active SLO protein reduced by TCEP (Sigma Aldrich) and 5 ug of corresponding IgG and/or IgG fragments. Xolair was purchased from Novartis. P.IgG represents a pool of immunoglobulin G isolated from the plasma of a donor who recently recovered from a GAS infection. A FragIT kit (Genovis) was utilized to digest P.IgG and generate two fractions: F(ab’)2- and Fc-fragments. Following incubation in ThermoMixer (Eppendorf) at 37°C and 300 rpm for 30 minutes, the plates were centrifuged, and the supernatant was transferred to a new plate for reading at 541 nm wavelength by a microplate reader (BMG Labtech), correlating to the quantity of leaked hemoglobin due to hemolysis. The positive control consisting of SLO alone was defined as a 100% lysis rate.

### *De novo* sequencing of the nAb, assembling protein and modeling of the antibody Fab

*De novo* sequencing and assembly of nAb was performed using BU-MS. The full-length nAb was initially reduced with 5 mM TCEP (Sigma Aldrich) and alkylated with 10 mM IAA (Sigma Aldrich), followed by overnight digestion in ThermoMixer (Eppendorf) at 37°C and 500 rpm, using trypsin, chymotrypsin, elastase, and pepsin (Promega) at an enzyme to substrate ratio of 1:20. The digested peptides were then purified using a C18 clean-up spin column (Thermo Scientific), concentrated in a SpeedVac (Eppendorf), and reconstituted into buffer A (2% acetonitrile, 0.2% formic acid) prior to mass spectrometry analysis.

Approximately 1 µg of peptides from each sample, quantified using a NanoDrop spectrophotometer (DeNovix), were loaded onto an EASY-nLC 1200 system interfaced with a Q Exactive HF-X hybrid quadrupole-Orbitrap mass spectrometer (Thermo Scientific). Each enzyme-digested sample was analyzed in duplicate injection. The peptides were first concentrated on a precolumn (PepMap100 C18 3 μm; 75 μm × 2 cm; Thermo Fisher Scientific) and then separated on an EASY-Spray column (ES903, column temperature 45 °C; Thermo Fisher Scientific), in according with the manufacturer recommendations. Two solvents were used as mobile phases: solvent A (0.1% formic acid) and solvent B (0.1% formic acid, 80% acetonitrile). A linear gradient from 5 to 38% B was employed over 180 minutes at a constant flow rate of 350 nl/min. For data acquisition, a data-dependent acquisition (DDA) method was implemented as follows. An initial MS1 scan with a scan range of 350–1650 m/z, resolution of 120,000, auto gain control (AGC) target of 3e6 and maximum IT (injection time) 45 ms was followed by the top 15 MS2 scans at a resolution of 15,000, AGC target 1e5, 30 ms IT and a stepped normalized collision energy (NCE) of 20, 25 and 30. Charge states of 1, 6-8 and above were excluded, except in samples digested with other enzymes than trypsin where singly charged ions were included. The performance of the LC-MS system was controlled by analyzing a yeast protein extract digest (Promega). A total of eight datasets were collected and subsequently processed using three de novo sequencing algorithms CasaNova, SuperNova and MEM (50).

A novel approach involving cumulative fragment-ion evidence was applied to enhance *de novo* peptide sequencing and subsequent primary protein structure assembly. Peptide candidates were generated using three deep-learning-based *de novo* peptide tools: PointNovo (51), CasaNovo (52), and InstaNovo (53). For PointNovo, two in-house multienzyme-trained models (54) were utilized, whereas default models were employed for CasaNovo and InstaNovo. The study considered twelve fragment ions: a+1, a+2, b+1, b+2, y+1, y+2, a-H2O, b-H2O, y-H2O, a-NH3, b-NH3, and y-NH3, with candidate selection based on a 20 ppm tolerance at both MS1 and MS2 levels and the observation of a minimum of four fragment ions. A positional evidence vector with length L was constructed for each candidate, filled by the count of found fragment ions at each position. These vectors were then pooled across all spectra from all samples.

Subsequently, peptides were segmented into 5-mers retaining the pooled positional fragment ion information. Cumulative MS evidence for these new 5-mer vectors was compiled elementwise. Finally, a cumulative MS score (cMS) for each peptide candidate was calculated by condensing the positional ion global evidence and normalizing for peptide length. This MS-evidence-driven strategy effectively disseminates information on redundant and overlapping segments identified across all MS samples analyzed. The highest-ranked peptide candidates were those that not only showed maximal overlap but also robust MS evidence supporting the target protein sequences. The positional confidence score (54) for the assembled heterodimeric chains of the Fab domain is illustrated in **Fig. 1F**.

Next, proABC-2 was employed to predict the hypervariable region of the nAb Fab domain, of relevance for subsequent docking studies (27). This was informed by the previously acquired sequences of both the heavy and light chains of the nAb. The structure of the Fab fragment of the nAb was predicted with AlphaFold-Multimer (v2.3.1) using the version-specific Docker container with default settings. The resulting structures were verified by calculating the inter-chain pDockQ values for each model (55). The predicted model with the highest pDockQ value was kept for downstream analysis.

### Crosslinking mass spectrometry and data analysis

The exploratory modeling process involved XL-MS for a crosslinking reaction and subsequent XL peptide identification. The nAb and SLO were mixed at a 1:1 molar ratio in 1xPBS solution at 37°C with 500 rpm agitation for an hour. The duplet linkers, DSS-H12/D12 or DSG-H6/D6 (Creative Molecules), were then added separately to crosslink the sample over the course of two hours. The reaction was quenched with 4 M ammonia bicarbonate (Sigma Aldrich), and followed by a standard reduction and alkylation procedures as stated above. We employed a two-step digestion process (involving lysyl endopeptidase, from FUJIFILM Wako Chemicals U.S.A. Corporation, followed by trypsin, Promega) in-solution to produce the crosslinked peptide pairs.

The peptides were cleaned, dried, and reconstituted before analysis by mass spectrometer as using the same protocol described above. Approximately 800 ng of the peptides from each sample, quantified using a NanoDrop spectrophotometer, were loaded into an Ultimate 3000 UPLC system connected to an Orbitrap Eclipse Tribrid Mass Spectrometer (Thermo Scientific). Each sample was performed technical duplicate injections. Column equilibration and sample loading were performed according to the manufacturer guidelines. The mobile phases used included Solvent A (0.1% formic acid) and Solvent B (0.1% formic acid and 80% acetonitrile). The gradient was linear and ranged from 5 to 38%, with a consistent flow rate of 300 nl/min over 90 minutes. The DDA method consisted of one MS1 scan with a scan range of 350– 1650 m/z, resolution of 120,000, a standard-mode AGC target and auto-mode maximum injection time. The fragment setup included a 3-second cycle time with MS2 scans, 15,000 resolution, standard AGC target, 22 ms IT and NCE of 30. All runs incorporated charge states from 2 to 6. The LC-MS performance was monitored prior to analysis using HeLa protein digest standard (Thermo Fisher Scientific).

The crosslinking datasets were analyzed using pLink2 (56), configuring DSG-H6/D6 and DSS-H12/D12 linker modifications according to the manufacturer product pages. The searched library included the sequences for the heavy, light chains of nAb and SLO along with common contaminants, and a maximum of two missed cleavage sites were allowed. Visualization of the top two nAb-SLO crosslinked peptides was generated using xiSPEC (57).

### DisVis interaction analysis based on distance constraints generated by XL-MS

All crosslinked peptides that connected the nAb Fab fragment and SLO, identified from both crosslinking experiments with duplet linkers, were extracted. DisVis was employed for quickscanning and interaction analysis (28), utilizing the crystal structure of SLO and the modeled Fab structure for exploratory modeling. Briefly, the distance between Cα-Cα derived from the interprotein crosslinked peptides was set within a range of 0-30 Å. Initially, all identified XL constraints were used to calculate the z-score and group the clusters. The cluster comprising four XLs, which included the two most abundant XLs, was further utilized in the interaction analysis. The prediction of accessible residues of both Fab and SLO was achieved using NetSurfP-3.0 (58), focusing on those with a relative solvent accessibility (RSA) greater than 40%. The interaction fraction index was computed through the interaction analysis function of DisVis (28), designating consistent IF values greater than 0.5 as putative residues that contribute to forming the suggested Fab-SLO binding interface. Above cut-off values were set according to DisVis developer manual to maximize confidence and reliability of the modeling practice.

### Distance-information-driven docking of Fab-SLO protein complex by HADDOCK

The HADDOCK 2.4 antibody-antigen information-driven docking protocol was implemented to construct the most accurate possible complex and to identify relevant interfacial residues (29, 31). Both the modeled Fab structure and the crystal structure of SLO were inputted, with the predicted HV loops of nAb assigned as active residues, and distance-derived putative interactive residues of SLO designated as passive residues. Two prevalent crosslink sites with a Cα-Cα range of 0-30Å were established as the center of mass constraints to enforce contact. Separate sampling parameters were set for rigid body docking, semi-flexible refinement, and final refinement at 10000, 400, and 400, respectively. From the candidate models, the complex with the lowest combined HADDOCK score was selected for downstream interface extraction following manual inspection. Finally, PRODIGY (32) was employed to classify the corresponding types of interaction within the interface, and SpotON (33) to predict hotspot residues, which are presumed to be considerably involved in intermolecular interactions.

### HDX-MS experiment and data analysis

The HDX-MS experimental setup involved a LEAP H/D-X PAL™ platform for automated sample preparation, which was interfaced with an LC-MS system that consisted of an Ultimate 3000 micro-LC connected to an Orbitrap Q Exactive Plus MS. HDX was carried out on SLO, both with and without commercially obtained nAb, in 10 mM PBS. Apo state (unbound SLO) and epitope mapping (Ab-bound SLO) samples were incubated for t = 0, 60, 1800, 9000 seconds at 20°C in either PBS or an HDX labelling buffer of identical composition prepared in D_2_O. The analysis was executed in a single, continuous run, with three replicates undertaken for each state and timepoint. The labelling reaction was quenched through dilution with 1% TFA, 0.4 M TCEP, 4 M urea, at 4°C. The quenched sample was directly injected and subjected to online pepsin digestion at 4°C. A flow of 50 μL/min 0.1% formic acid was applied for 4 minutes for online digestion and trapping of the samples. Digestion products underwent online solid phase extraction and washing with 0.1% FA for 60s on a trap column (PepMap300 C18), which was switched in-line with a reversed-phase analytical column (Hypersil GOLD). Separation occurred at 1°C, with mobile phases of 0.1 % formic acid (A) and 95% acetonitrile/0.1% formic acid (B), using a gradient of 5-50% B over 8 minutes and then from 50 to 90% B for 5 minutes. The separated peptides were analyzed on a Q Exactive Plus MS, equipped with a heated electrospray source (HESI) operating at a capillary temperature of 250°C with sheath gas 12 au, auxiliary gas 2 au, and sweep gas 1 au. For HDX analysis, MS full scan spectra were obtained at 70,000 resolution, automatic gain control (AGC) 3e6, Max ion injection time 200ms and scan range 300-2000 m/z. The identification of generated peptides was performed by analyzing separate un-deuterated samples using data dependent MS/MS. A library pool of peptides that included peptide sequence, charge state, and retention time was created for the HDX analysis by running pepsin-digested, un-deuterated samples against the SLO sequence on PEAKS Studio X (Bioinformatics Solutions Inc.). HDX data analysis and visualization were performed using HDExaminer v3.1.1 (Sierra Analytics Inc.).

Ab-bound states were analyzed compared to Apo states, using a single charge state per peptide. Given the comparative nature of the measurements, the deuterium incorporation for the peptic peptides was derived from the observed relative mass difference between the deuterated and non-deuterated peptides without back-exchange correction using a fully deuterated sample. The spectra for all time points were manually inspected; low scoring peptides, obvious outliers, and peptides for which retention time correction could not be made consistent were removed. Deuteros 2.0 (59, 60) was further applied to perform the hybrid significance test and visualization the change of deuterium uptake in coverage plot, butterfly plot, volcano plot, kinetic uptake, as well as projecting the protected peptide residue coordinates to the indicated 3D structure.

### Carriage and epitope conserveness analysis, and reverse-engineering of the construct

A dedicated BLAST database was established using genomic data from Streptococcus pyogenes sourced from The Bacterial and Viral Bioinformatics Resource Center (BV-BRC) (35) as of March 28, 2023. Genomes classified as being of poor quality, sourced from plasmids, or as duplicate entries were excluded. This led to the creation of a curated database comprising 2216 genomes, narrowed down from the original 2283 entries. Employing BLASTp (61), the Streptolysin O sequence (Uniprot ID: P0DF96) was queried against this database. Hits that covered over 98% of the query sequence were considered matches, with the threshold selected based on data characteristics, as detailed in the supplementary material. Domain 3 of Streptolysin O from Streptococcus pyogenes M3, spanning residues 250-299 and 346-420, was identified as the target for construct design. This domain, interrupted by a subsequence of Domain 1, required a re-engineered approach for continuity. The Message Passing Neural Network ProteinMPNN (62) algorithm was utilized to determine an alternate amino acid sequence capable of maintaining the original backbone conformation of Domain 3. The redesign process commenced with the substitution of the Domain 1 sequence with a tetra-glycine linker, chosen for its structural flexibility. This modified sequence’s quaternary structure was predicted using AlphaFold2 (34) and subsequently aligned to the native structure in ChimeraX to confirm that Domain 3’s integrity was preserved post-substitution. Ensuring the structural stability of the designed construct was paramount. An assessment was conducted to ascertain if substituting any internal amino acids could yield a more stable conformation. ProteinMPNN was employed to evaluate potential replacements for the internal residues, while all surface residues remained unchanged. This optimization process revealed that the native internal residues were optimal, and no substitutions were necessary.

The workflow illustration was created by BioRender.com. GraphPad Prism 10 was used to generate dot plots, bar lots, and heatmaps, and perform statistics and regression analysis. ChimeraX and Chimera (63) were applied to visualize corresponding crosslinking pseudo-bonds, to depict 3D structures of proteins, domains, and epitopes, and to perform superimposition and alignment.

## Acknowledgement

We gratefully acknowledge the Swedish National Infrastructure for Biological Mass Spectrometry (BioMS), the SciLifeLab Integrated Structural Biology platform, and the Protein Production Sweden (PPS) for providing facilities and experimental support. We would also like to thank Dr. Hong Yan, Dr. Anahita Bakochi, Dr. Wolfgang Knecht and Dr. Mikael Lindberg for assistance.

## References

1. S. Boyoglu-Barnum, et al., Quadrivalent influenza nanoparticle vaccines induce broad protection. Nature 592, 623–628 (2021).

2. J. M. Steichen, et al., A generalized HIV vaccine design strategy for priming of broadly neutralizing antibody responses. Science 366 (2019).

3. A. Lanzavecchia, A. Frühwirth, L. Perez, D. Corti, Antibody-guided vaccine design: identification of protective epitopes. Curr Opin Immunol 41, 62–67 (2016).

4. M. I. Anasir, C. L. Poh, Structural Vaccinology for Viral Vaccine Design. Front Microbiol 10, 738 (2019).

5. I. E. Mba, et al., Vaccine development for bacterial pathogens: Advances, challenges and prospects. Trop Med Int Health 28, 275–299 (2023).

6. P. Pushparaj, et al., Immunoglobulin germline gene polymorphisms influence the function of SARS-CoV-2 neutralizing antibodies. Immunity 56, 193–206.e7 (2023).

7. L. Hanke, et al., Multivariate mining of an alpaca immune repertoire identifies potent cross-neutralizing SARS-CoV-2 nanobodies. Sci Adv 8, eabm0220 (2022).

8. S. Ständer, L. R. Grauslund, M. Scarselli, N. Norais, K. Rand, Epitope Mapping of Polyclonal Antibodies by Hydrogen–Deuterium Exchange Mass Spectrometry (HDX-MS). Anal Chem 93, 11669–11678 (2021).

9. S. Chowdhury, et al., Dissecting the properties of circulating IgG against Group A Streptococcus through a combined systems antigenomics-serology workflow. *bioRxiv*, 2023.11.07.565977 (2023).

10. O. Ozohanics, A. Ambrus, Hydrogen-Deuterium Exchange Mass Spectrometry: A Novel Structural Biology Approach to Structure, Dynamics and Interactions of Proteins and Their Complexes. Life 10, 286 (2020).

11. C. Yu, L. Huang, Cross-Linking Mass Spectrometry: An Emerging Technology for Interactomics and Structural Biology. Anal Chem 90, 144–165 (2018).

12. J. Mintseris, S. P. Gygi, High-density chemical cross-linking for modeling protein interactions. Proc National Acad Sci 117, 93–102 (2020).

13. S. Hauri, et al., Rapid determination of quaternary protein structures in complex biological samples. Nat Commun 10, 192 (2019).

14. W. Bahnan, et al., A human monoclonal antibody bivalently binding two different epitopes in streptococcal M protein mediates immune function. Embo Mol Med (2022) 10.15252/emmm.202216208.

15. S. Lenz, et al., Reliable identification of protein-protein interactions by crosslinking mass spectrometry. Nat Commun 12, 3564 (2021).

16. N. J. Avire, H. Whiley, K. Ross, A Review of Streptococcus pyogenes: Public Health Risk Factors, Prevention and Control. Pathogens 10, 248 (2021).

17. E. Chiarot, et al., Targeted Amino Acid Substitutions Impair Streptolysin O Toxicity and Group A Streptococcus Virulence. Mbio 4, e00387–12 (2013).

18. K. van Pee, et al., CryoEM structures of membrane pore and prepore complex reveal cytolytic mechanism of Pneumolysin. Elife 6, e23644 (2017).

19. S. Uchiyama, et al., Streptolysin O Rapidly Impairs Neutrophil Oxidative Burst and Antibacterial Responses to Group A Streptococcus. Front Immunol 6, 581 (2015).

20. A. Azuar, et al., Recent Advances in the Development of Peptide Vaccines and Their Delivery Systems against Group A Streptococcus. Nato Adv Sci Inst Se 7, 58 (2019).

21. B. Limbago, V. Penumalli, B. Weinrick, J. R. Scott, Role of Streptolysin O in a Mouse Model of Invasive Group A Streptococcal Disease. Infect Immun 68, 6384–6390 (2000).

22. M. R. Davies, et al., Atlas of group A streptococcal vaccine candidates compiled using large-scale comparative genomics. Nat Genet 51, 1035–1043 (2019).

23. D. R. Walkinshaw, et al., The Streptococcus pyogenes vaccine landscape. Npj Vaccines 8, 16 (2023).

24. A. J. Farrand, S. LaChapelle, E. M. Hotze, A. E. Johnson, R. K. Tweten, Only two amino acids are essential for cytolytic toxin recognition of cholesterol at the membrane surface. Proc National Acad Sci 107, 4341–4346 (2010).

25. I. Kucinskaite-Kodze, M. Simanavicius, J. Dapkunas, M. Pleckaityte, A. Zvirbliene, Mapping of Recognition Sites of Monoclonal Antibodies Responsible for the Inhibition of Pneumolysin Functional Activity. Biomol 10, 1009 (2020).

26. R. Evans, et al., Protein complex prediction with AlphaFold-Multimer. Biorxiv, 2021.10.04.463034 (2022).

27. F. Ambrosetti, et al., proABC-2: PRediction Of AntiBody Contacts v2 and its application to information-driven docking. Bioinformatics 36, btaa644 (2020).

28. G. C. P. van Zundert, et al., The DisVis and PowerFit Web Servers: Explorative and Integrative Modeling of Biomolecular Complexes. J Mol Biol 429, 399–407 (2017).

29. F. Ambrosetti, B. Jiménez-García, J. Roel-Touris, A. M. J. J. Bonvin, Modeling Antibody-Antigen Complexes by Information-Driven Docking. Structure 28, 119–129.e2 (2020).

30. F. Ambrosetti, Z. Jandova, A. M. J. J. Bonvin, Computer-Aided Antibody Design. Methods Mol Biology 2552, 267–282 (2022).

31. S. J. de Vries, M. van Dijk, A. M. J. J. Bonvin, The HADDOCK web server for data-driven biomolecular docking. Nat Protoc 5, 883–897 (2010).

32. L. C. Xue, J. P. Rodrigues, P. L. Kastritis, A. M. Bonvin, A. Vangone, PRODIGY: a web server for predicting the binding affinity of protein–protein complexes. Bioinformatics 32, 3676–3678 (2016).

33. I. S. Moreira, et al., SpotOn: High Accuracy Identification of Protein-Protein Interface Hot-Spots. Sci. Reports 7, 8007 (2017).

34. J. Jumper, et al., Highly accurate protein structure prediction with AlphaFold. Nature 596, 583–589 (2021).

35. R. D. Olson, et al., Introducing the Bacterial and Viral Bioinformatics Resource Center (BV-BRC): a resource combining PATRIC, IRD and ViPR. Nucleic Acids Res. 51, D678– D689 (2022).

36. G. Pantaleo, B. Correia, C. Fenwick, V. S. Joo, L. Perez, Antibodies to combat viral infections: development strategies and progress. Nat. Rev. Drug Discov. 21, 676–696 (2022).

37. A. W. Senior, et al., Improved protein structure prediction using potentials from deep learning. Nature 577, 706–710 (2020).

38. J. A. Ruffolo, L.-S. Chu, S. P. Mahajan, J. J. Gray, Fast, accurate antibody structure prediction from deep learning on massive set of natural antibodies. Nat Commun 14, 2389 (2023).

39. R. Yin, B. Y. Feng, A. Varshney, B. G. Pierce, Benchmarking AlphaFold for protein complex modeling reveals accuracy determinants. Protein Sci 31, e4379 (2022).

40. J. T. Seffernick, S. Lindert, Hybrid methods for combined experimental and computational determination of protein structure. J. Chem. Phys. 153, 240901 (2020).

41. E. V. Petrotchenko, C. H. Borchers, Protein Chemistry Combined with Mass Spectrometry for Protein Structure Determination. Chem Rev 122, 7488–7499 (2022).

42. Z. Orbán-Németh, et al., Structural prediction of protein models using distance restraints derived from cross-linking mass spectrometry data. Nat Protoc 13, 478–494 (2018).

43. R. Jia, R. T. Bradshaw, V. Calvaresi, A. Politis, Integrating Hydrogen Deuterium Exchange–Mass Spectrometry with Molecular Simulations Enables Quantification of the Conformational Populations of the Sugar Transporter XylE. J. Am. Chem. Soc. 145, 7768– 7779 (2023).

44. G. R. Masson, M. L. Jenkins, J. E. Burke, An overview of hydrogen deuterium exchange mass spectrometry (HDX-MS) in drug discovery. Expert Opin Drug Dis 12, 981–994 (2017).

45. E. D. Merkley, et al., Distance restraints from crosslinking mass spectrometry: Mining a molecular dynamics simulation database to evaluate lysine–lysine distances. Protein Sci 23, 747–759 (2014).

46. M. M. Zhang, et al., An Integrated Approach for Determining a Protein–Protein Binding Interface in Solution and an Evaluation of Hydrogen–Deuterium Exchange Kinetics for Adjudicating Candidate Docking Models. Anal. Chem. 91, 15709–15717 (2019).

47. K. A. Manthei, et al., Structural analysis of lecithin:cholesterol acyltransferase bound to high density lipoprotein particles. *Commun*. Biol. 3, 28 (2020).

48. F. Hugo, J. Reichwein, M. Arvand, S. Krämer, S. Bhakdi, Use of a monoclonal antibody to determine the mode of transmembrane pore formation by streptolysin O. Infect. Immun. 54, 641–645 (1986).

49. S. C. Feil, D. B. Ascher, M. J. Kuiper, R. K. Tweten, M. W. Parker, Structural Studies of Streptococcus pyogenes Streptolysin O Provide Insights into the Early Steps of Membrane Penetration. J Mol Biol 426, 785–792 (2014).

50. D. Beslic, G. Tscheuschner, B. Y. Renard, M. G. Weller, T. Muth, Comprehensive evaluation of peptide de novo sequencing tools for monoclonal antibody assembly. Brief Bioinform 24, bbac542 (2022).

51. R. Qiao, et al., Computationally instrument-resolution-independent de novo peptide sequencing for high-resolution devices. *Nat*. Mach. Intell. 3, 420–425 (2021).

52. M. Yilmaz, et al., Sequence-to-sequence translation from mass spectra to peptides with a transformer model. *bioRxiv*, 2023.01.03.522621 (2023).

53. K. Eloff, et al., De novo peptide sequencing with InstaNovo: Accurate, database-free peptide identification for large scale proteomics experiments. bioRxiv, 2023.08.30.555055 (2023).

54. C. Gueto-Tettay, et al., Multienzyme deep learning models improve peptide de novo sequencing by mass spectrometry proteomics. Plos Comput Biol 19, e1010457 (2023).

55. P. Bryant, G. Pozzati, A. Elofsson, Improved prediction of protein-protein interactions using AlphaFold2. Nat. Commun. 13, 1265 (2022).

56. Z.-L. Chen, et al., A high-speed search engine pLink 2 with systematic evaluation for proteome-scale identification of cross-linked peptides. Nat Commun 10, 3404 (2019).

57. L. Kolbowski, C. Combe, J. Rappsilber, xiSPEC: web-based visualization, analysis and sharing of proteomics data. Nucleic Acids Res. 46, gky353-(2018).

58. M. H. Høie, et al., NetSurfP-3.0: accurate and fast prediction of protein structural features by protein language models and deep learning. Nucleic Acids Res 50, gkac439-(2022).

59. A. M. Lau, J. Claesen, K. Hansen, A. Politis, Deuteros 2.0: Peptide-level significance testing of data from hydrogen deuterium exchange mass spectrometry. Bioinformatics 37, btaa677-(2020).

60. A. M. C. Lau, Z. Ahdash, C. Martens, A. Politis, Deuteros: software for rapid analysis and visualization of data from differential hydrogen deuterium exchange-mass spectrometry. Bioinformatics 35, btz022-(2019).

61. C. Camacho, et al., BLAST+: architecture and applications. BMC Bioinform. 10, 421 (2009).

62. J. Dauparas, et al., Robust deep learning–based protein sequence design using ProteinMPNN. Science 378, 49–56 (2022).

63. E. F. Pettersen, et al., UCSF Chimera—A visualization system for exploratory research and analysis. J. Comput. Chem. 25, 1605–1612 (2004).

